# Extracellular DNA traps in a ctenophore demonstrate conserved immune cell behaviors in a non-bilaterian

**DOI:** 10.1101/2020.06.09.141010

**Authors:** Lauren E. Vandepas, Caroline Stefani, Phillip P. Domeier, Nikki Traylor-Knowles, Frederick W. Goetz, William E. Browne, Adam Lacy-Hulbert

**Author notes:** Equal contribution (Lauren E. Vandepas, Caroline Stefani).

## Abstract

The formation of extracellular DNA traps (ETosis) is a first response mechanism by specific immune cells following exposure to microbes ^1,2^. Initially characterized in vertebrate neutrophils, cells capable of ETosis have been discovered recently in diverse non-vertebrate taxa ^3 4-6^. To assess the conservation of ETosis between evolutionarily distant non-vertebrate phyla, we observed and quantified ETosis using the model ctenophore *Mnemiopsis leidyi* and the oyster *Crassostrea gigas*. Here we report that ctenophores – thought to have diverged very early from the metazoan stem lineage ^7-10^– possess immune cell types capable of phagocytosis and ETosis. We demonstrate that both *Mnemiopsis* and *Crassostrea* immune cells undergo ETosis after exposure to diverse microbes and chemical agents that stimulate ion flux. Our results support ETosis as an evolutionarily ancient metazoan defense against pathogens.

## Main

Response to invading microbes is a primary physiological task for all metazoans. In vertebrates, specialized immune cell types such as macrophages and neutrophils detect the presence of microbes. These immune cells perform specific behaviors like phagocytosis or secretion of antimicrobial compounds to sequester and eliminate microbial invaders ^11,12^. The release of extracellular DNA traps (ETs) is a relatively recently described immune cell behavior – a morphologically and molecularly distinct cell death process called ETosis – during which immune cells cast filamentous nets composed of nuclear chromatin material or mitochondrial DNA into the surrounding extracellular space, trapping and killing invading microbes ^13,14^. Initially believed to be a behavior exclusive to vertebrate neutrophils, recent studies have highlighted ETosis as an anti-microbial response in non-vertebrate taxa ^4,5,15-18^. The independent evolution of ETs has also been proposed in taxa as divergent as social amoebas ^19^ and plants ^20^.

In mammalian cells, ETosis and other immune responses are often initiated by pattern recognition receptors (PRRs) that bind to molecular motifs on microbes commonly referred to as pathogen-associated molecular patterns (PAMPs) ^21^. PRR protein sequences and domain architectures can vary between taxa, as they evolve to detect host specific pathogens ^22-24^. For example, mussels have a radiation of TLRs that may recognize a suite of specific pathogens ^25^, while *Drosophila* uses its Toll receptors to identify the protein Spätzle ^26^. Despite these broad patterns of clade-specific immune receptor diversity, the signaling cascades and effectors downstream of PRRs have been shown to be well conserved ^30^, including intracellular signaling pathways involving calcium, MAP kinases, and reactive oxygen species (ROS) ^27-29^.

In vertebrate leukocytes, exposure to cytokines, microbes, PAMPs, or pharmacological agents activate specific cellular pathways involved in ET formation, such as a response to ion imbalance (calcium, potassium) or reactive oxygen species (ROS) burst ^31,32^. Although the detailed mechanisms of these pathways remain unknown in non-vertebrates ^5,6,14,31,33^, genes involved in vertebrate immune cell signaling cascades have been identified in diverse metazoans, including non-bilaterians ^34-38^.

Non-vertebrate ETosis has been best studied in molluscan model systems, particularly in bivalves like the model oyster *Crassostrea gigas* ^4-6,16,39,40^. However, attempts to stimulate production of ETs in *Crassostrea* via exposure to some microbial signatures as well as ETosis-stimulating drugs has yielded variable results ^6^. For example, robust ETotic response in oyster hemocytes was observed following challenge with *Vibrio* ^4^, however induction of ETosis with other microbes or PAMPs has been either modest or not observed ^6^. Without a consensus assessment of ETosis stimulation, the conservation of ET induction in non-vertebrates remains unclear, and it is unknown whether ETosis represents a fundamental metazoan immune response ^4-6^.

Ctenophores, also known as “comb jellies”, are a clade of gelatinous planktonic marine animals that diverged early from the animal stem lineage and represent one of the most ancient extant metazoan phyla ^8-10^. Ctenophores have two distinct germ layers (ectoderm and endomesoderm) separated by a jelly-like layer of collagenous mesoglea and lack a circulatory system. Understanding functional attributes of ctenophore physiological systems has provided fundamental insights into the conservation of animal cell types and signaling pathways, as well as revealing mechanisms for the emergence of evolutionary novelties ^36,41-43^. Currently, the ctenophore immune system remains almost entirely undescribed ^36,44^. Specific immune cell types have not been explicitly identified in ctenophores, though they possess motile amoebocyte-like cells that are abundant in the mesoglea and are capable of phagocytosis ^36,45^. Whether ctenophores have additional specialized cellular immune mechanisms has not been reported.

Here we demonstrate that the model ctenophore *Mnemiopsis leidyi* possesses immune cell types capable of ETosis in response to diverse stimuli known to activate conserved signaling cascades in vertebrate immune cells. To rapidly and accurately quantify ETosis we developed a novel, automated imaging pipeline to identify ETs. We applied this imaging pipeline to comparatively assess ET production in *Mnemiopsis* and the bivalve mollusk *Crassostrea gigas*, a model for invertebrate ETosis. Our data from both non-vertebrate species suggest that cells undergoing ETosis following a range of stimuli represent evolutionarily ancient anti-microbial defense mechanisms in metazoans.

## Results

### Isolated ctenophore cells display immune behaviors

To investigate whether the model ctenophore *Mnemiopsis leidyi* possesses cell types with specialized immune functions, we mechanically disassociated whole *Mnemiopsis* and examined isolated cells ^46,47^. We observed morphologically distinct motile cell types displaying prominent intracellular granules and/or vesicles, including amoebocyte-like cells with relatively short pseudopodia and stellate cells with long pseudopodia (**Fig. 1A-C, Supp. Video 1**). Time-lapse microscopy shows that some stellate cells have a dynamic morphology and can rapidly vary the length and number of pseudopodia (**Supp. Video 2)**. These motile cells are highly active in primary cell cultures and their morphologies and scavenging behaviors are reminiscent of immune cell types, such as macrophage-like cells, that have been described in diverse metazoans ^11^.

**Figure 1.**
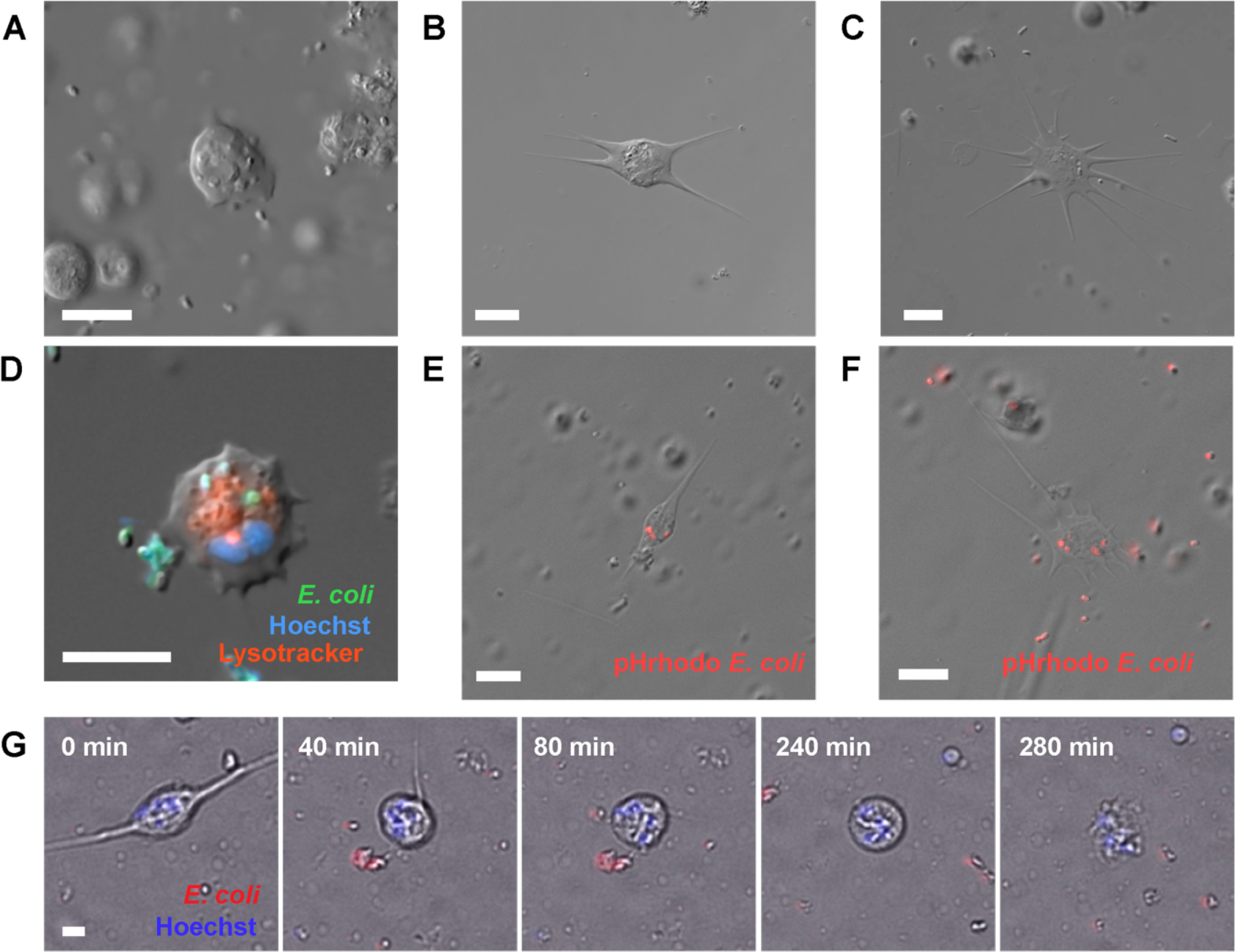
Stellate and amoebocyte-like *Mnemiopsis* cells display immune behaviors. (A) DIC image of a highly granular amoebocyte-like cell. (B) DIC image of a granular stellate cell displaying multiple processes. (C) DIC image of a stellate cell with pseudopodia. (D) Merged brightfield and fluorescent image of a live *Mnemiopsis* motile, stellate cell that has phagocytosed fluorescent *E. coli* (green). Nuclei are labeled with Hoechst (blue). Lysosomes are labeled with Lysotracker-redDND99 (red). Scale bar is 10 μm. (E)-(F) Combined DIC and fluorescent images of live granular cells with pseudopodia that have phagocytosed fluorescent *E. coli*. (G) Still images from Movie S2 showing a motile stellate cell retracting its processes, undergoing nuclear rotation (spinning), and extruding its nuclear contents after exposure to *E. coli*.

A major role for macrophage-like cells is host defense against microbes. We therefore assessed whether the motile cells present in *Mnemiopsis* exhibited fundamental immune cell behaviors in response to bacteria. *Mnemiopsis* cells were incubated with killed fluorescently labeled *Escherichia coli* and imaged by combined DIC and fluorescence microscopy. We found *Mnemiopsis* cells exhibiting diverse morphologies internalized fluorescent *E. coli* ^36^; these included motile stellate and amoeboid cells (**Fig. 1D-F**). Furthermore, timelapse DIC microscopy revealed that amoeboid cells actively maneuvered their pseudopodia to engulf nearby bacteria, supporting a potential role in sensing and removing microbes (**Supp. Video 3**).

Following exposure to *E. coli* a conspicuous subset of stellate cells changed their morphology dramatically by retracting their pseudopodial processes and undergoing nuclear rotation, followed by a rapid expulsion of cellular material (**Fig. 1G; Supp. Video 4; Supp. Video 5**). This behavior is remarkably similar to vertebrate monocyte behavior preceding extracellular DNA trap (ET) formation ^31,48^. This led us to speculate that some ctenophore immune cells were producing ETs in response to the presence of microbes.

### Ctenophore immune cells produce extracellular traps composed of chromatin

In vertebrates, ETs are released in response to recognition of microbial components, and protect against infection by immobilizing and killing invading microbes^1^. To test whether incubation with microbes promoted extrusion of DNA from *Mnemiopsis* cells we stained isolated cells with Hoechst to label DNA. In isolated cells incubated in seawater media, we observed intact nuclei with concentrated Hoechst labeling (**Fig. 2A**). After treatment with heat-killed fluorescent *E. coli* we observed networks of Hoechst-stained DNA filaments cast in large areas around some individual *Mnemiopsis* cell bodies. Three-dimensional rendering of confocal z-stacks showed that *Mnemiopsis* cells extruding DNA had bacteria entangled in the DNA networks, consistent with ETs (**Fig. 2B; Supp. Video 6;** ^49^).

**Figure 2.**
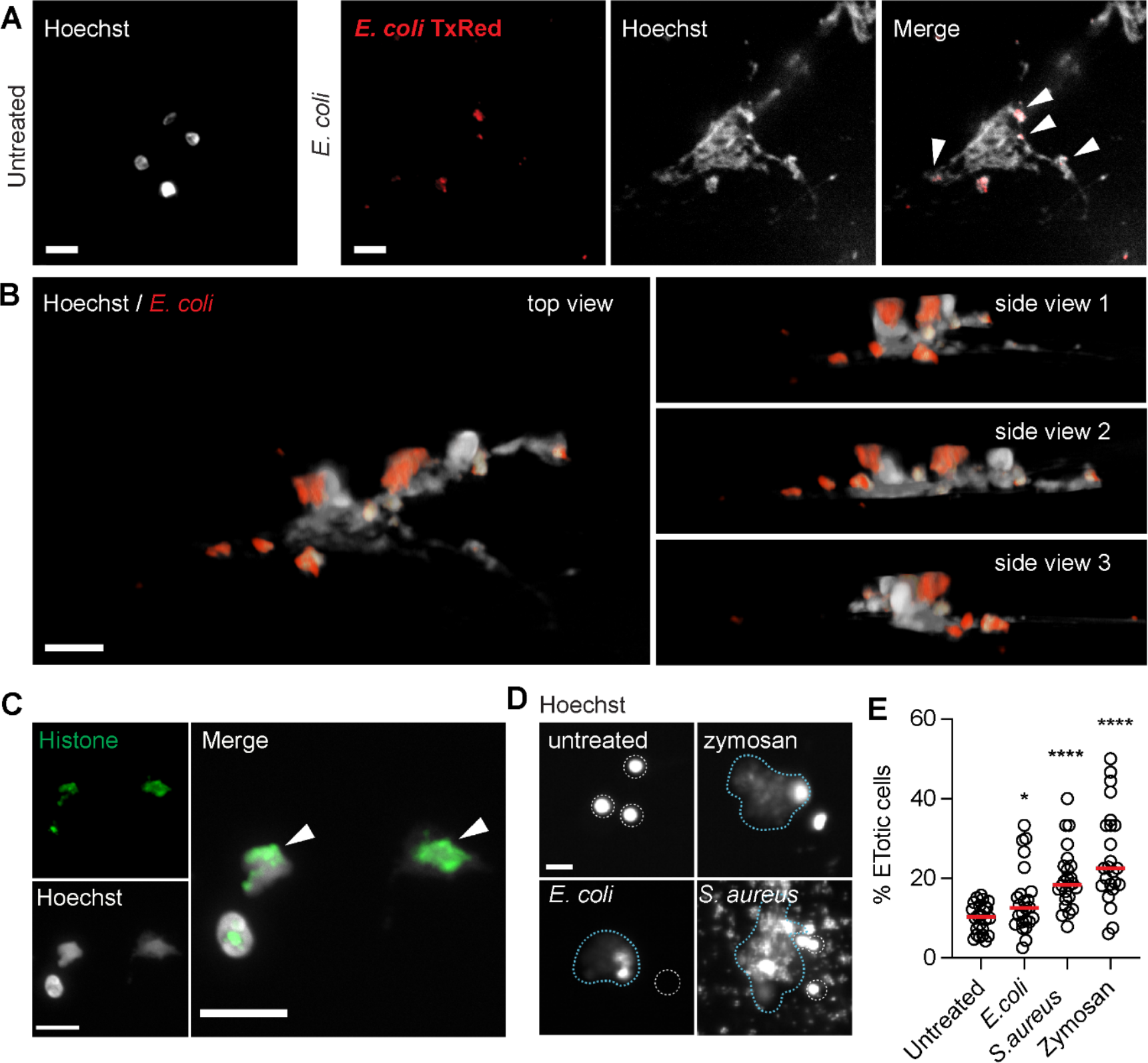
Ctenophore immune cells undergo ETosis when exposed to microbes. (A)-(C) Confocal images of *Mnemiopsis* cell nuclei stained with Hoechst. (A) Left – Nuclei of unstimulated *Mnemiopsis* cells. Right – A cell exposed to fluorescent TxRed-*E. coli* has undergone ETosis; a large web-like pattern of chromatin has been extruded from the cell. Individual *E. coli* can be seen enmeshed by chromatin filaments (white arrowheads). (B) Still images from a 3D projection of *Mnemiopsis* ET. (C) Confocal image of *Mnemiopsis* extracellular traps composed of DNA and histones. Histone 11-4 antibody (green) and Hoechst (white) staining are visible in intact and ETosed *Mnemiopsis* cells treated with the potassium ionophore nigericin. White arrowhead marks extracellular DNA+histone chromatin nets. (D) Fluorescent microscope (Cytation) images of Hoechst-labeled *Mnemiopsis* cells. Representative images of intact nuclei are outlined in white dotted lines, while boundaries of ETs are marked with blue dotted lines. (E) Incubation with *S. aureus, E. coli*, and zymosan significantly induced ETosis in *Mnemiopsis* cells. Scale bar is 10 μm in all images.

In vertebrate immune cells, ETs can originate either from the cell nucleus or from mitochondria ^50,51^. To assess the organellar origin of the extruded DNA, we stained *Mnemiopsis* cells with an antibody that recognizes an array of nuclear histone proteins (H1, H2A, H2B, H3, H4). Nuclei of non-ETotic *Mnemiopsis* cells were labeled with this histone antibody, which showed stereotypical patterns of concentrated DNA and histone labeling, confirming the specificity of this antibody in *Mnemiopsis* cells (**Fig. 2C**) ^52,53^. Notably, in cells that have undergone ETosis, the extracellular ET DNA also stained with this pan-histone antibody, demonstrating that the extruded DNA filaments are composed of chromatin (**Fig. 2C, arrows**). Together, these data identify that *Mnemiopsis* cells produce *bona fide* extracellular DNA traps and further concluded that these ETs are derived from the nucleus ^14^.

### Ctenophore ETosis is an immune response to microbial exposure

To assess whether the formation of ETs in *Mnemiopsis* is a response to microbial challenge, we measured ET production after incubation of *Mnemiopsis* cells with *E. coli* and two additional microbial particles: heat-killed gram-positive *S. aureus*, and the yeast cell wall extract zymosan. We found that all three microbial particles stimulated ET formation in *Mnemiopsis* cells when compared with untreated cells incubated with media alone (**Fig. 2D**). Numbers of ETotic cells were manually quantified based on release of Hoechst-positive material by fluorescent microscopy (**Fig. 2E**). Exposure to *S. aureus* and zymosan promoted considerably higher rates of ET formation (mean increase of 98.2% and 117.7% over control respectively) than *E. coli* (43.7%) (**Fig. 2E**). These data confirm that production of ETs represents a *Mnemiopsis* immune cell-specific response to the presence of microbes.

### Ctenophore ETs form filamentous networks that capture microbes

Scanning electron microscopy (SEM) is commonly used to confirm the structure of ETs, and is informative in distinguishing ETosis from DNA release secondary to other forms of cell death such as necrosis or apoptosis ^54^. To examine the structure of the extruded DNA in *Mnemiopsis* cell cultures, we performed SEM on isolated cells that had been incubated with *S. aureus* or with seawater media alone. Images of untreated *Mnemiopsis* cells confirm the presence of diverse morphological cell types observed by DIC, including stellate cells (**Fig. 3A; Supp. Fig. 1)**. Following incubation with *S. aureus*, we observed long filamentous networks surrounding some *Mnemiopsis* cells (**Fig. 3B-C; Supp. Fig. 1**), which are similar to ETs produced by vertebrate and crustacean immune cells ^1,5,55,56^. We also observed a close association between individual *S. aureus* and the filamentous nets produced by *Mnemiopsis* cells, providing further evidence that these structures are able to capture microbes, a critical function of ETs **(Fig. 3B-C insets**) ^57^.

**Figure 3.**
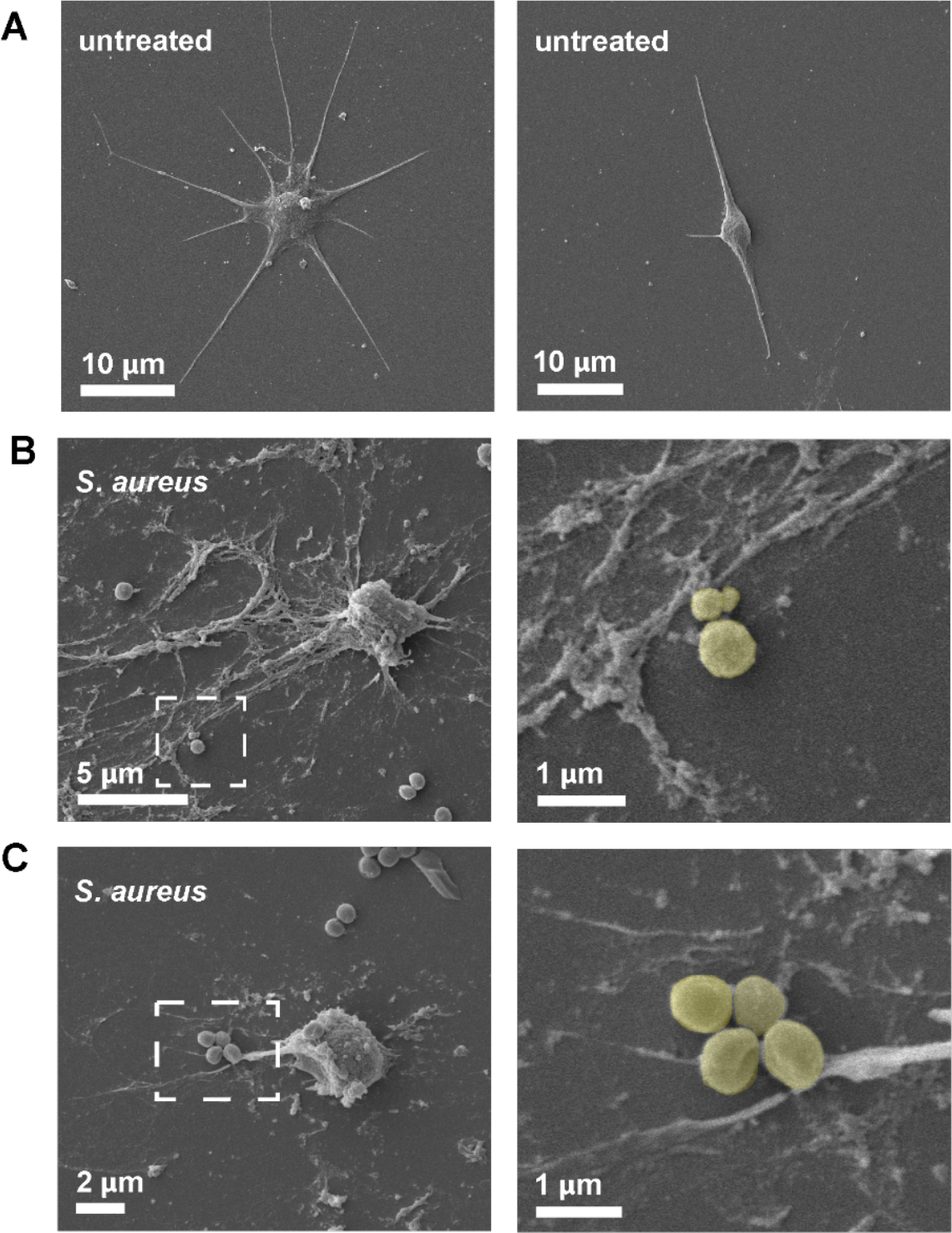
SEM of *Mnemiopsis* ETs. Scanning electron microscopy (SEM) images of isolated *Mnemiopsis* cells. (A) Images of untreated *Mnemiopsis* stellate cells showing multiple pseudopodial processes. (B)-(C) *Mnemiopsis* cells incubated with *S. aureus* produce extracellular traps composed of long filamentous networks surrounding the cells. Individual *S. aureus* bacterium can be seen closely associated with the filaments. Insets show high magnification of *S. aureus* closely associated with the *Mnemiopsis* ETs. (*S. aureus* are pseudocolored in high magnification for clarity).

### PMA induces ETosis in *Mnemiopsis*

We hypothesized that ETosis in ctenophores may be initiated by engaging evolutionarily conserved signaling pathways known to precede ETosis in other metazoans. For example, phorbol 12-myristate 13-acetate (PMA) is commonly used to induce ET production in vertebrate immune cells through activation of NADPH-mediated ROS ^31^. This pathway has been proposed to induce ET formation in immune cells in annelids and crustaceans ^5,15^, suggesting that it may be evolutionarily ancient and well conserved in metazoans. We treated *Mnemiopsis* cells with PMA for four hours, stained DNA with Hoechst, and visualized cells by fluorescence microscopy. We observed the extrusion of DNA after PMA treatment, which was similar ot that seen after micobial challenge (**Fig. 2D; Fig. 4A**).

**Figure 4.**
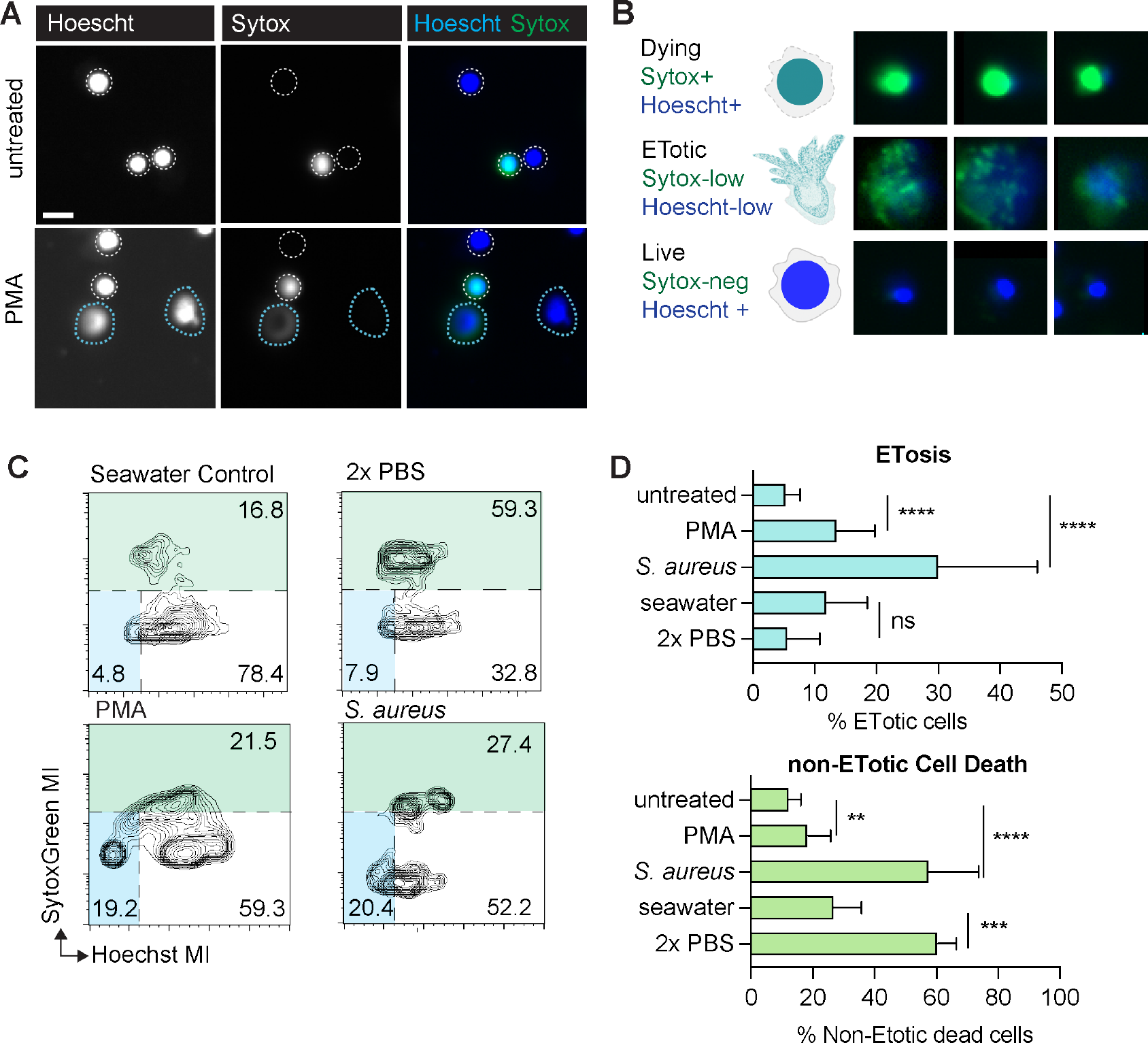
PMA stimulation in *Mnemiopsis* evaluated by an automated imaging analysis pipeline. All samples were incubated with their respective stimuli or seawater media alone for 4 hours and imaged. (A) Representative images of untreated controls (top) and PMA-incubated (bottom) *Mnemiopsis* cells labeled with Hoechst and Sytox Green. White dotted lines denote rough boundaries of intact nuclei. Blue dotted lines outline ETs. (B) Representative images of nuclei from SytoxGreen^high^ cells which are dead or dying non-ETotic cells (top), Hoechst^low^/ SytoxGreen^low^ ETotic cells showing diffuse Hoechst and SytoxGreen fluorescence (middle), Hoechst^high^/ SytoxGreen^low^ live cells that show intact cell membranes. (C) Representative FlowJo plot showing distribution of fluorescent signals from *Mnemiopsis* cells labeled with Hoechst and SytoxGreen following respective treatments. (D) ETosis is significantly stimulated in *Mnemiopsis* cells are incubated with PMA gram-positive *S. aureus in vitro*. Total cell death increases significantly after incubation with dilute saline compared to seawater control, while detection of ETosis events is unchanged.

Importantly, exposure to PMA can also trigger non-ETotic cell death pathways ^58^, raising the possibility that the extracellular DNA we observed was a result of cell rupture and release of DNA secondary to cell death. To distinguish between these possible scenarios, we used a combination of two DNA dyes: Hoechst, which is a membrane-permeable dye that labels DNA in both live cells and dead cells, and SytoxGreen, a non-permeable cell stain that selectively labels DNA in cells with compromised cell membranes, as observed in cells undergoing cell death (**Fig. 4A**). Both DNA dyes label ETs ^5,59^, and ETotic cells have both diffuse gradients of Hoechst and Sytox staining (**Fig. 4A-B**). We then developed a semi-automated imaging analysis approach that allowed us to measure Hoescht and SytoxGreen labeling in individual cell nuclei and accurately quantify both ETosis and cell death simultaneously (**Fig. 4B; Supp. Fig. 2**). ET production has been successfully identified and quantified in neutrophils using a combination of machine learning and high-throughput imaging ^60,61^, we deployed similar techniques to assess ETosis in ctenophore cells. An approach similar to those implemented in flow cytometry analyses was used in which we plotted the intensity of SytoxGreen fluorescence against Hoechst signal for each individual cell (**Fig. 4B-C**). This allowed for the identification of three distinct populations in PMA-treated *Mnemiopsis* cells: Hoechst^high^/ SytoxGreen^low^ live cells that show no DNA release and intact cell membranes; Hoechst^low^/ SytoxGreen^low^ cells which display diffuse extracellular Hoechst and SytoxGreen fluorescence, representing ETotic cells ^5,59^; and SytoxGreen^high^ cells with a range of Hoescht staining, which are dead or dying non-ETotic cells.

To confirm that the image analysis pipeline accurately distinguishes non-ETotic cell death from ETosis, we induced death in *Mnemiopsis* cells by exposure to a moderate osmotic shock (dilute saline solution, 2X PBS). We measured a significant increase in non-ETotic dead cells compared with normal seawater media controls (**Fig. 4C**). Notably, dying cells were clearly separated from cells undergoing ETosis. Using this pipeline, we then quantified ETosis and cell death in cells treated with PMA. Confirming our initial observations by microscopy, PMA promoted significant induction of ETosis, with a mean increase of 57.2% over cells incubated with seawater media alone (**Fig. 4C-D**). Importantly, non-ETotic cell death did not significantly increase following PMA incubation.

We sought to establish that the automated image analysis pipeline could detect ETosis induced by microbial exposure. Because we observed clear ETs with SEM following *S. aureus* challenge, we then applied this automated imaging approach to this microbial stimulus. Samples that are incubated with microbes contain more background fluorescence due to Hoescht and Sytox labeling of *S. aureus* nuclear material; however, we observed a clear increase in ETosis following *S. aureus* challenge using automated imaging analysis, as well as an increase in non-ETotic cell death (**Fig. 4C and 4D**). Together, these data confirm that PMA induces ETosis in *Mnemiopsis*. Further, the novel semi-automated image analysis pipeline presented here can accurately detect and quantify invertebrate ETosis and distinguish ETosis from total cell death.

### ETosis in *Mnemiopsis* is initiated by agonists that induce ion flux

In vertebrates, ET formation can be triggered by membrane ion flux triggered by exposure to bacterial toxins that act as ionophores ^31,53,62^. Because stimulation of ETosis via ion flux has limited reports in non-vertebrates ^6,32^, we assessed whether well-characterized bacterially-derived ionophores nigericin (potassium ionophore) and calcimycin (A23187; calcium ionophore) could also elicit ETosis in *Mnemiopsis* cells. We incubated *Mnemiopsis* cells with the K^+^ ionophore nigericin and observed a mean increase of 367% over untreated cells (**Fig. 5A and 5B; Supp. Video 6; Supp. Fig. 3**). Incubation with the Ca^2+^ ionophore calcimycin resulted in a mean increase of 225% over untreated control cells (**Fig. 5A and 5B)**. Furthermore, although incubation with either ionophore also significantly increased *Mnemiopsis* cell death, consistent with their effects in vertebrate cells ^63-65^, our image analysis pipeline was able to clearly distinguish ETosis from other forms of cell death. These data therefore support an evolutionarily conserved pathway for ET formation in response to immune stimulation or cellular damage.

**Figure 5.**
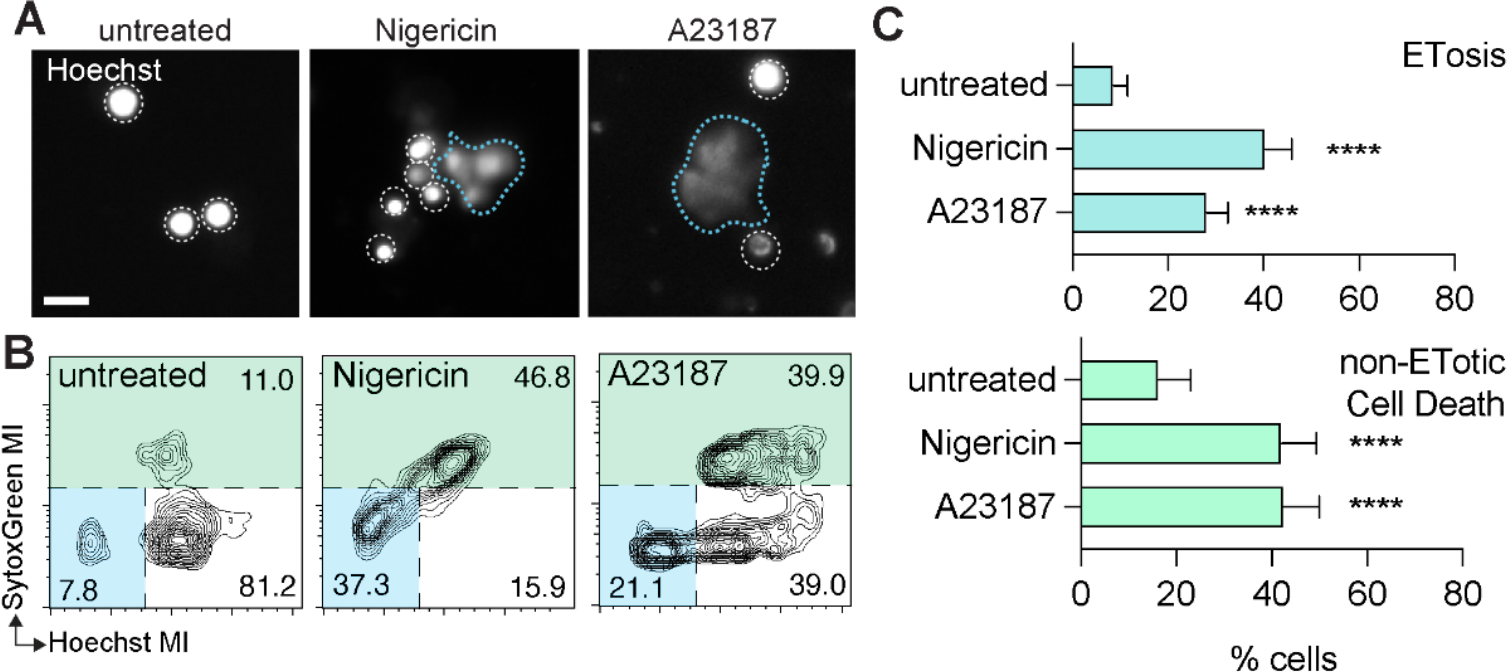
ETosis in *Mnemiopsis* is initiated by agonists that induce ion flux. (A) Representative Hoechst-labeled nuclei of untreated controls (left) and cells incubated with either K^+^ ionophore nigericin or Ca^2+^ ionophore A23187. White dotted lines denote rough boundaries of intact nuclei. Blue dotted lines outline ETs. (B) Representative FlowJo plot showing distribution of fluorescent signals from cells labeled with Hoechst and SytoxGreen labeled following respective treatments. (C) ETosis is significantly stimulated in *Mnemiopsis* cells incubated with K^+^ ionophore nigericin and Ca^2+^ ionophore A23187. Total cell death also increases significantly after incubation with both ionophores.

### Microbes, PMA, and calcium ionophore stimulates ETosis in *Crassostrea gigas* hemocytes

Our observations that ET formation in *Mnemiopsis* can be triggered by similar signaling pathways to those involved in mouse and human immune cell ETosis supports the hypothesis that ETosis represents an evolutionarily ancient response to infection or cell damage activated by intracellular signaling pathways that are widely conserved throughout animals ^66-68^. However, stimulation of ETosis has not been widely assessed outside of vertebrates ^6,32^ or compared between phyla. To compare stimulation of *Mnemiopsis* cells with another non-vertebrate metazoan that is an established model for analysis of ETosis, we extended our analysis to the bivalve mollusc, *Crassostrea gigas*, a marine non-vertebrate bilaterian with a circulatory system possessing specialized immune cells (hemocytes). Some hemocyte cell types have been shown to display immune behaviors such as phagocytosis ^69^. We incubated isolated hemocytes with fluorescent *E. coli* for 4 hours to assess whether *C. gigas* cells were responding to the presence of bacteria by phagocytosing the *E. coli* (**Fig. 6A-B)** or undergoing ETosis (**Fig. 6B**, inset). When observing hemocytes by confocal microscopy, we found that some hemocytes are highly phagocytic, with individual cells ingesting multiple bacteria (**Fig. 6B)**. Fluorescent imaging also identified networks of extracellular DNA surrounding some hemocytes after exposure to microbial particles, similar to previous reports of bivalve ET formation ^4^ (**Fig. 6B, insets**).

**Figure 6.**
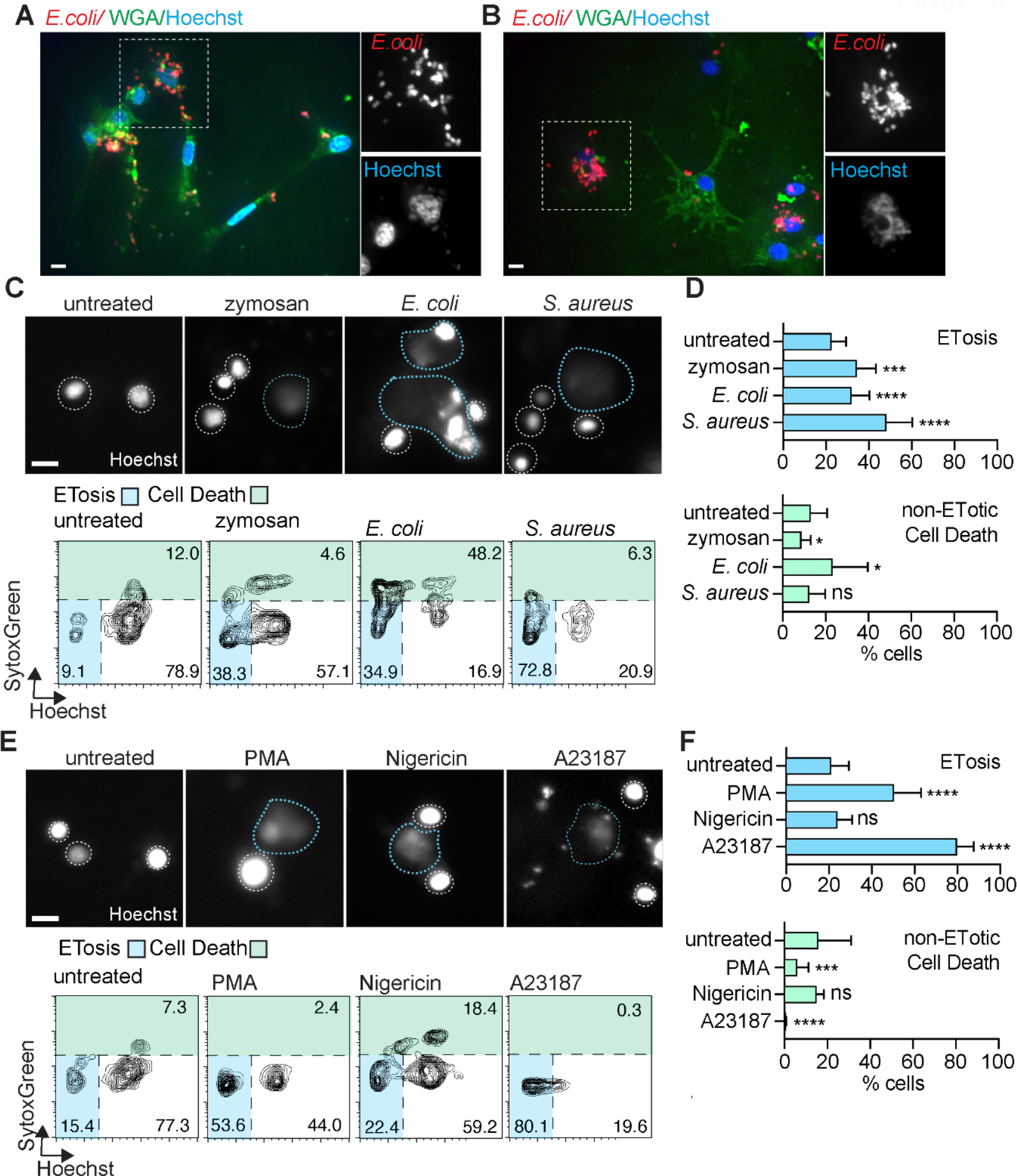
Diverse stimuli induce ETosis in Crassostrea gigas hemocytes. (A) Confocal image showing *C. gigas* hemocytes that have phagocytosed fluorescent *E. coli*. (B) Confocal image with *C. gigas* hemocytes where some have phagocytosed fluorescent bacteria (arrow) and one cell has produced an ET (arrowhead). Insets show *E. coli* and diffuse DNA staining indicative of ETosis of a hemocyte. (C) (top) Representative images of hemocyte nuclei stained with Hoechst that were either incubated with media alone or exposed to fungal or bacterial PAMPs. White dotted lines denote rough boundaries of intact nuclei. Blue dotted lines outline ETs. (bottom) Representative FlowJo plot showing distribution of fluorescent signals from *C. gigas* hemocytes labeled with Hoechst and SytoxGreen following respective treatments. (D) ETosis is significantly stimulated in *Crassostrea* hemocytes exposed to zymosan, gram-negative *E. coli*, and gram-positive *S. aureus* compared to seawater media controls after 4 hours of incubation. Non-ETotic cell death decreases slightly following zymosan incubation, increases slightly with *E. coli* treatment. No significant change in non-ETotic cell death is detected with *S. aureus* treatment.(E) (top) Representative images of hemocyte nuclei stained with Hoechst that were either incubated with media alone or with PMA, K^+^ ionophore nigericin, or Ca^2+^ ionophore A23187. White dotted lines denote rough boundaries of intact nuclei. Blue dotted lines outline ETs. (bottom) Representative FlowJo plot showing distribution of fluorescent signals from *C. gigas* hemocytes labeled with Hoechst and SytoxGreen following respective treatments. (F) PMA and A23187, but not nigericin, significantly stimulate ETosis in hemocytes. Observations of non-ETotic cell death in hemocytes treated with PMA or A23187 decrease. Scale bar is 10 μm in all images.

While ETs have been reported in multiple bivalve species, successful induction of ETosis in hemocytes following incubation with either microbial stimuli or pharmaceuticals has varied ^4-6^. To directly compare ET production in oyster hemocytes with observations in *Mnemiopsis*, we assessed ETosis in *C. gigas* hemocytes using the same panel of microbes. We first confirmed that *Crassostrea* immune cells undergo ETosis in response to microbial stimulation using the automated imaging analysis pipeline (**Fig. 6C-D**). Hemocytes were incubated with heat-killed *E. coli, S. aureus*, or zymosan, and then assessed using the automated image analysis pipeline for quantification of ETosis. We found that ETosis increased after exposure to all three microbes, when compared to hemocytes incubated *in vitro* with seawater media alone (**Fig 6C-D)**. *Crassostrea* hemocytes undergo ETosis in response to bacterial challenge with both gram-negative *E. coli* and gram-positive *S. aureus* with mean increases of 52.3% and 41.5% over untreated hemocytes, respectively (**Fig. 6D)**. The fungal PAMP zymosan also elicited a significant ETotic response in oyster hemocytes, with a mean increase of 112% over seawater media controls. Interestingly, *S. aureus* stimulation did not stimulate significant increases in non-ETotic cell death in hemocytes (**Fig. 6D**), in contrast to what we observed in *Mnemiopsis* cells (**Fig. 2D-E**). Hemocyte total non-ETotic cell death increased significantly with either *E. coli* or zymosan exposure.

We also measured ETosis in *Crassostrea* hemocytes after four-hour incubations with PMA, nigericin, calcimycin, or seawater media controls (**Fig. 6E-F**). Exposure to PMA induced significant ET production in isolated *Crassostrea* hemocytes, with a relative mean increase of 95.7% (**Fig. 6F**). A large proportion of *Crassostrea* hemocytes also produced ETs after exposure to calcimycin, with a mean increase of 209% over controls (**Fig. 6F**). These results are similar to the robust response reported in prior studies of bivalve hemocytes to Ca^2+^ ionophore stimulation ^6^. In contrast to *Mnemiopsis* cells, incubation of hemocytes with the K^+^ ionophore nigericin showed no significant induction of ETosis (compare **Fig. 2D-E and Fig. 6F**). Notably, non-ETotic cell death does not increase in *C. gigas* hemocytes following drug incubation. Unexpectedly, both PMA and calcimycin treatment resulted in a relatively minor, but significant, decrease in non-ETotic cell death.

## Discussion

Identifying the core molecular mechanisms of immune cells and their evolutionary origins is critical for understanding the evolution of metazoan immune cell function and behavior. Here we report that ctenophores, which diverged very early from the metazoan stem lineage, have immune cells capable of ETosis, a cellular response to microbes first described in mammalinan neutrophils. We also show that cells isolated from the model ctenophore *Mnemiopsis leidyi* undergo ETosis following exposure to diverse microbial components, and stimulation of intracellular signaling cascades known to precede ETosis in vertebrates. For example, we find that incubation with the potassium ionophore nigericin stimulates ETosis in *Mnemiopsis* cells, as has been reported for mammalian neutrophils, suggesting that K^+^ efflux may be an evolutionarily conserved trigger for ETosis. We conclude from our results that ETosis represents an ancient mechanism of cellular immune defense in metazoans (**Fig. 7**).

**Figure 7.**
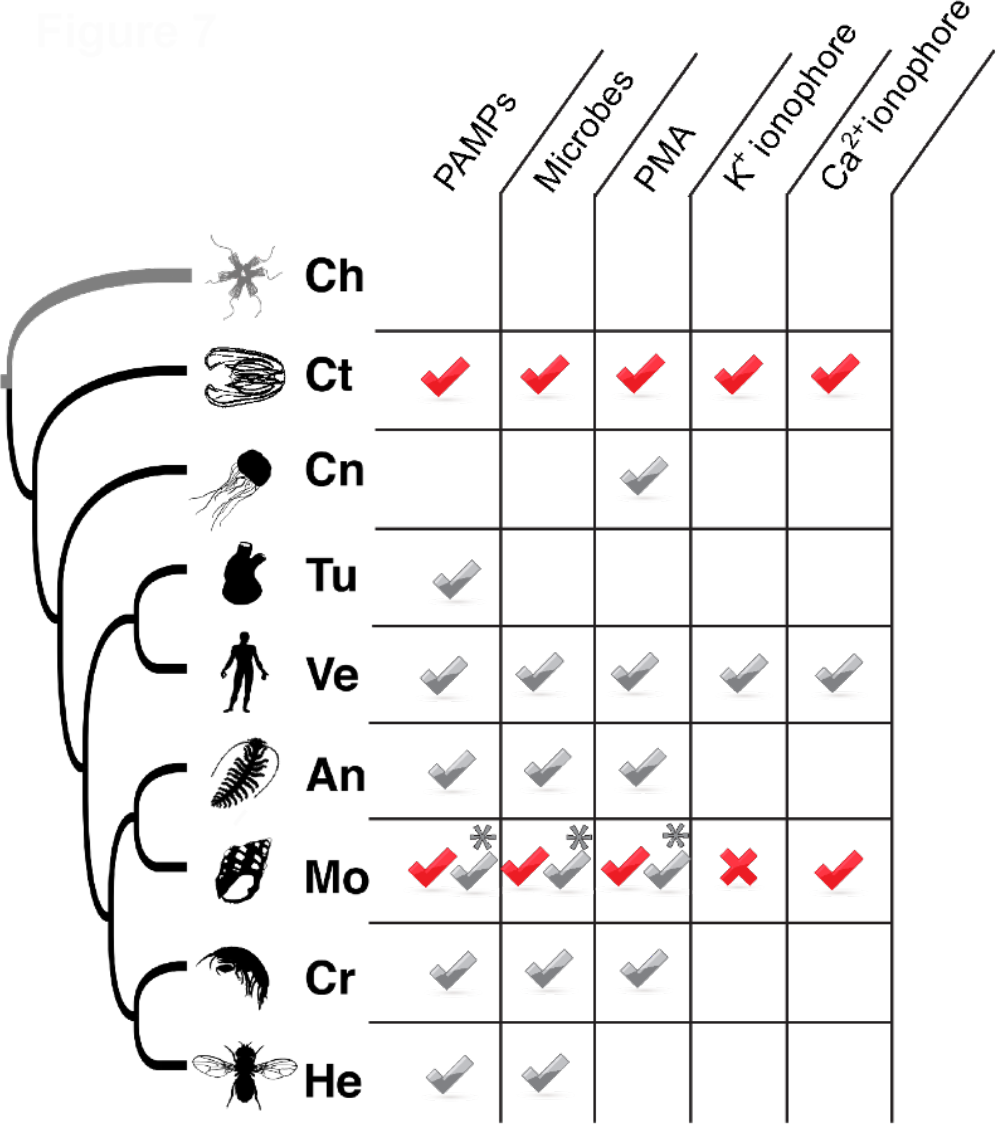
ETosis is an ancient metazoan immune response. Summary of ETosis phenomena across Metazoa. The presence of cells competent for ETosis in diverse non-bilaterian, protostome, and deuterostome taxa support the production of extracellular DNA traps as an ancient animal immune defense mechanism. Taxa abbreviations: Ch – Choanoflagellata, Ct – Ctenophora, Cn – Cnidaria, Tu – Tunicata, Ve – Vertebrata, An – Annelida, Mo – Mollusca, Cr – Crustacea, He – Hexapoda. Red – this study. Grey – other studies. * – Conflicting reports of ETosis stimulation.

Since the first description of neutrophil extracellular traps ^11^, multiple pathways involved in the generation of mammalian ETs have been identified ^31^. However, the intracellular actors necessary for ET formation in non-vertebrate taxa are less well understood. For example, gene homologs for proteins known to be essential for ETosis in mammalian neutrophils, such as peptidylarginine deiminase 4 (PAD4), neutrophil elastase, myeloperoxidase (MPO) and pannexin-1 are absent outside of vertebrates. There is strong evidence for the presence of multiple ETosis pathways in non-vertebrates that can be activated via stimulation with microbial signatures, parasites, PMA, A23187, and UV light ^4-6,15-17^. Potassium ion efflux, for example, is classically associated with cellular damage, including during bacterial infection ^70^. Calcium flux has been implicated in ET formation in vertebrates either as a component of immune receptor-mediated signaling or as an activator of downstream effectors that drive DNA condensation preceding ETosis ^71^. In addition to intracellular ion flux, components of vertebrate ETosis pathways, such as ERK, p38, or Akt, are foundational components of metazoan intracellular signaling ^66-68^.

Our ability to make further direct comparisons of the molecular mechanisms of ETosis between vertebrates and *Mnemiopsis* is currently hampered by a lack of tractable forward genetic approaches and is therefore largely restricted to pharmacological perturbation. However, our data supports considerable apparent conservation of ET formation; firstly, bacterial and fungal components (*E. coli*, intact *S. aureus* and fungal particles) significantly stimulated ETosis in *Mnemiopsis* and *Crassostrea* hemocytes in our study. Our results are in agreement with work in other non-vertebrates and supports the concept that ET formation is a fundamental component of metazoan anti-microbial defense ^4-6,15-17^. We also demonstrate that ETosis in *Mnemiopsis* and *Crassostrea* is stimulated by both PMA, which in mammalian cells stimulates protein kinase C and Ca^2+^ flux, and by the Ca^2+^ ionophore A23187. These findings align with observations in vertebrate neutrophils ^31^ supporting an evolutionarily conserved role for Ca^2+^ transients in ETosis. Finally, the K^+^ ionophore nigericin promoted robust ETosis in *Mnemiopsis*, indicating additional conservation of K^+^ sensing and signaling pathways. Interestingly, *Crassostrea* hemocytes did not produce ETs in response to nigericin, raising the possibility that the nigericin-stimulated pathway is lost in this mollusc species. Signaling pathway inhibitors optimized for use in marine invertebrate cells will be useful for future studies of molecular actors involved in immune responses.

In contrast to vertebrate immune cell nuclear morphologies that correspond with discrete immune cell functions (e.g. lobed nuclei in neutrophils and eosinophils; ^72^), no obvious differences in nuclear morphologies have been reported in either bivalve hemocytes or ctenophore immune cells. We did observe that *Mnemiopsis* immune cells that underwent ETosis *in vitro* after exposure to *E. coli* did not appear to have phagocytosed bacteria in significant amounts (**Supp. Video 2**). Intriguingly, we also observed other motile cells that had phagocytosed large amounts of bacteria without undergoing ETosis (**Supp. Fig. 2, Supp. Video 4; Supp. Video 5**). Our functional characterization of ETosis-competent *Mnemiopsis* cells suggests that future studies should assess whether ETosis-competent and highly phagocytic non-ETotic cells represent discrete immune cell types in ctenophores.

Ctenophores have not had specific immune cell antimicrobial behaviors described beyond phagocytosis ^36,45^. We demonstrate that *Mnemiopsis leidyi* possesses cells functionally competent for ETosis in response to a range of microbial challenges. It remains unclear whether non-vertebrate immune cells are capable of vital (mitochondrial) ETosis, which involves production of ETs derived from mitochondrial DNA while maintaining cell viability ^18^. Our data demonstrates that both *Mnemiopsis leidyi* and *Crassostrea gigas* immune cells can be stimulated to produce ETs with distinct pharmaceutical agents that, in vertebrate immune cells, differentially induce specific intracellular signaling pathways to activate ET production. The production of extracellular traps in *Mnemiopsis* suggests this specific immune cell antimicrobial defense behavior was likely present early in metazoan evolution, prior to the diversification of extant metazoan lineages.

## Methods

### Animal maintenance

Laboratory cultures of *Mnemiopsis leidyi* were maintained as previously described (Presnell, 2021 #35). Adult *Mnemiopsis* cells were isolated from whole animals following established protocols ^36,46,47^. *Crassostrea gigas* were maintained under flowing seawater at approximately 13°C and hemolymph was extracted from the adductor muscle with a syringe. Isolated cells from both taxa were maintained *in vitro* under sterile conditions in 0.22μm filtered seawater (FSW) + 1% penicillin/streptomycin.

### Scanning electron microscopy (SEM)

*Mnemiopsis* cells were seeded onto coverslips treated with poly-L-lysine. Cells were incubated with or without heat-killed *E. coli* for 4 hours to induce ET formation. Cells were fixed in ½ strength Karnovsky’s fixative (2.5% glutaraldehyde, 2% paraformaldehyde in 0.1M sodium cacodylate buffer, pH 7.3) overnight at 4°C. Samples were then rinsed with 0.1M cacodylate buffer and treated with 1% osmium tetroxide for 1 hour. The samples were then dehydrated through a graded series of alcohols and critical point dried (Autosamdri, Tousimis Corp, Rockville, MD). Samples were mounted on stubs, sputter coated with gold/palladium (Denton Desk IV, Denton Vacuum, Moorestown, NJ) and imaged on a JSM 6610 LV scanning electron microscope at 5kV (JEOL, Tokyo, Japan).

### Stimulation of ETosis and imaging

For stimulation and quantification of ETosis, cells were isolated from 24 individual animals, plated in 96 well plates, and exposed to pHrodo-*E. coli, Staphylococcus aureus*, zymosan particles (Sigma Aldrich), 25 uM nigericin (Thermo Fisher Scientific), 1 mg/mL PMA (Sigma Aldrich), or 4 uM A23187 (Sigma). Each experimental condition for each animal was performed in triplicate. Live cell staining was performed following (2) and live imaging was performed using a JuLI Stage (NanoEntek).

For immunofluorescence, ctenophore cells were prepared as previously described ^46,47,73^ and labeled with mouse anti-histone H11-4 (EMD Millipore) and anti-mouse Alexa Fluor 488 (Thermo Fisher Scientific). The H11-4 histone antibody was selected because it recognizes histones H1, H2A/B, H3, and H4 proteins across diverse species. Cells were imaged using ×60 objective and ×100 oil objective, on a Nikon Ti (Eclipse) inverted microscope with Ultraview Spinning Disc (CSU-X1) confocal scanner (Perkin Elmer). Images were captured with an Orca-ER Camera using Volocity (Quorum technologies). Post-acquisition analysis such as contrast adjustment, deconvolution through iterative restoration and colocalization were performed using Volocity software.

For quantification of ETosis and total cell death, cells were treated with Hoechst 33342 (Sigma Aldrich) and SytoxGreen (Invitrogen) for 20 min, before imaging at 20X on an automated imaging plate reader, Cytation 3 (Biotek, software Gen5 v4.2).

### Automated image-based profiling

We analyzed over 28,000 total images using CellProfiler (v4.1.3). Image quality was assessed by calculating a focus score using two classes Otsu thresholding method, weighted variance on 20×20 pixel measurements. We calculated and applied an illumination correction for each fluorescent channel (SytoxGreen and Hoechst) using a background illumination function of 50 pixels block size, without smoothing. Each corrected image was then segmented using a global robust background method (0.05-50), with a smoothing scale of 1.3488 and a correction factor of 0.89. Clumped objects were identified and split by shape. For each segmented object we measured the number and intensity of pixels in each fluorescent channel. Each image and each segmented object, along with Metadata, were exported as csv files by experiment. R (v4.0.5) software with tidyverse (v1.3.1), dplyr (v1.0.7) and readr (v1.4.0) packages were then used to transform the datasets. Data from images and objects were merged, and measurements from individual images with a Focus Score <0.2 were removed from further analysis. This allowed us to identify and select only images that were in focus. Surface area, Hoechst intensity and SytoxGreen intensity per object (nucleus) and per individual animal were then imported into FlowJo (v10.8.0), and percentages of cells per delineated population (dead/dying cell, live cell, and ETotic cell) were calculated. Dying and ETotic cells were gated as indicated on the figures. Finally, percentages per individual animal surveyed were combined and tested for statistical significance using GraphPad Prism (v9.2.0). All statistical tests were performed using two-tailed unpaired student t-test *p<0.05, **p<0.01, ***p<0.001, ****p<0.0001.

## Acknowledgements

The authors are grateful to Anna Bruchez and Rachel Prins for imaging support. This research was supported by the National Oceanographic and Atmospheric Administration, a National Research Council Postdoctoral Fellowship to LEV, and the National Science Foundation under Grant No. 2013692. CS and ALH were supported by National Institutes of Health Grants R33AI119341 and R01GM102482. The authors thank Steve Ziegler for technical and experimental support. The authors gratefully acknowledge the Fred Hutchinson Cancer Center Electron Microscopy Shared Resource (EMSR). The EMSR is supported in part by the Cancer Center Support Grant P30 CA015704-40.

## Author contributions

Conceptualization, C.S. and L.E.V.; Methodology, L.E.V. and C.S.; Formal Analysis, C.S. and L.E.V.; Investigation, L.E.V. and C.S.; Visualization, C.S., L.E.V., and W.E.B.; Writing – Original Draft, L.E.V. and C.S.; Writing – Review & Editing, C.S., A.L.H., W.E.B., F.W.G., and L.E.V.; Funding Acquisition, A.L.H., W.E.B., N.T.K., F.W.G., and L.E.V.; Resources, A.L.H., W.E.B., and F.W.G.; Supervision, A.L.H., W.E.B., and F.W.G.

## Supplemental Material

**Supp. Figure 1:** Widefield views of SEM images of unstimulated (left panel) and microbe-challenged (right panel) *Mnemiopsis* cells *in vitro*. Unstimulated cells from whole *Mnemiopsis* show diverse sizes and morphologies indicating the presence of multiple cell types. Microbe-exposed *Mnemiopsis* cells were incubated with *S. aureus* for 3 hours.

**Supp. Figure 2:** Workflow schematic of the automatic imaging analysis pipeline.

**Supp. Video 1:** Video of DIC timelapse images of isolated *Mnemiopsis* cells *in vitro*. An amoebocyte-like granular cell (arrowhead) and highly motile stellate cell (arrow) are visible.

**Supp. Video 2**: Video of DIC timelapse images of an isolated *Mnemiopsis* stellate cell *in vitro*. The cell initially has two large processes, then absorbs them and subsequently produces several additional processes. The cell then begins to crawl out of view.

**Supp. Video 3:** Merged brightfield and fluorescent video of a live *Mnemiopsis* motile, stellate cell that is phagocytosing fluorescent *E. coli* (green). *Mnemiopsis* cells were labeled with Lysotracker Red. Scale bar is 10 μm.

**Supp. Video 4**: Merged brightfield and fluorescent video of a live *Mnemiopsis* motile, stellate cell undergoing ETosis after *in vitro* exposure to pHrodo-*E. coli*. The cell moves into view, retracts its processes, spins, and exudes its nuclear material. DNA (Hoechst, blue), pHrodo*-E. coli* (red)

**Supp. Video 5:** 3D reconstruction of confocal stack of *Mnemiopsis* extracellular DNA traps. DNA (Hoechst, white) is extracellular, in filamentous “nets”; lysosomal marker (Lysotracker, green) denotes cellular debris; bacteria are ensnared in the “nets” (pHrodo-*E. coli*, red)

**Supp. Video 6:** Merged brightfield and fluorescent video of a live *Mnemiopsis* motile, stellate cell that is phagocytosing fluorescent *E. coli* (red). Scale bar is 10 μm.

