## Supplementary material for "Extracellular DNA traps in a ctenophore demonstrate conserved immune cell behaviors in a non-bilaterian": Supplementary Material.pdf

[a]NRC Research Associateship Program; [b]Northwest Fisheries Science Center, National Oceanographic and Atmospheric Administration, Seattle, WA 98112; [c]Benaroya Research Institute at Virginia Mason, Seattle, WA 98101; [d]University of Miami Rosenstiel School of Marine and Atmospheric Sciences, Miami, FL 33149; [e]University of Miami Department of Biology, Coral Gables, FL 33146; [†]Corresponding author; *Equal contribution

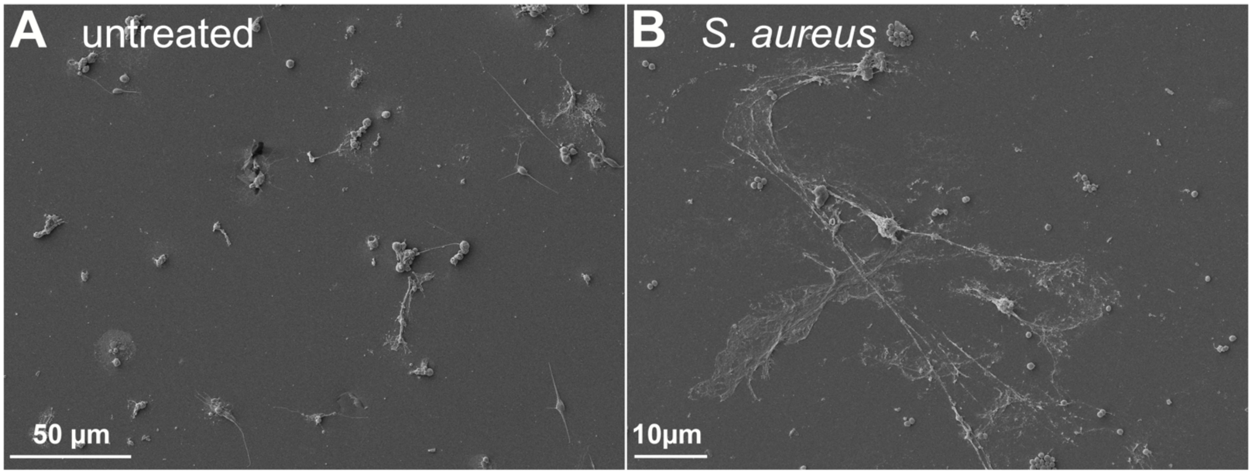


**Supp. Figure 1: Widefield views of SEM images of unstimulated (left panel) and microbe-challenged (right panel) *Mnemiopsis* cells *in vitro*.** Unstimulated cells from whole *Mnemiopsis* show diverse sizes and morphologies indicating the presence of multiple cell types. Microbe-exposed *Mnemiopsis* cells were incubated with *S. aureus* for 3 hours.

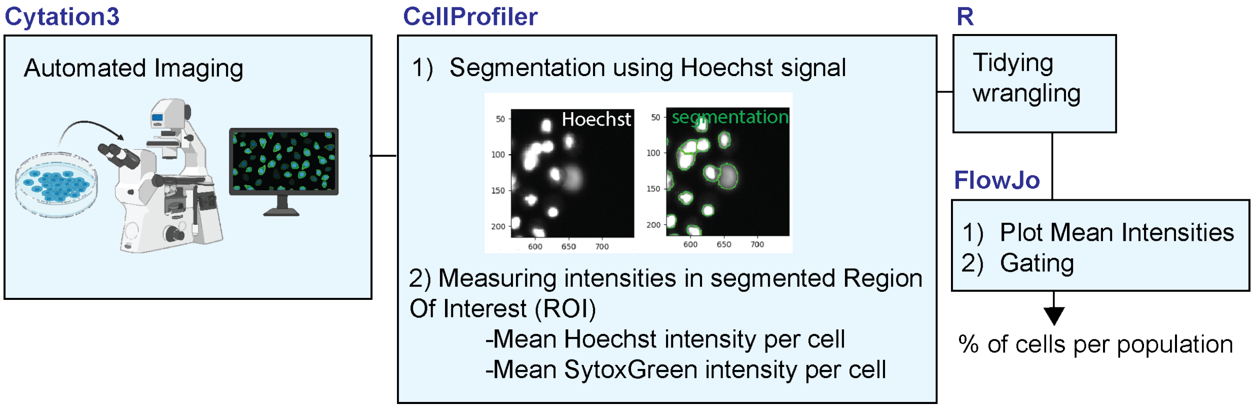


**Supp. Figure 2: Workflow schematic of the automatic imaging analysis pipeline**. We imaged cells using CellProfiler. Image quality was measured and the background was corrected prior to segmentation. We measured the mean intensity of pixels in each fluorescent channel inside the segmented Region Of Interest (ROI). Each measurement along with Metadata, were exported as csv files. Datasets were then tidied and wrangled in R, to prepare them for import into FlowJo. Hoechst intensity and SytoxGreen intensity per object (nucleus) and per individual animal were then imported into FlowJo, and percentages of cells per delineated population (dead/dying cell, live cell, and ETotic cell) were calculated. Dying and ETotic cells were gated as indicated in figure 4.

**LEGEND SUPPLEMENTARY VIDEOS:**

**Supp. Video 1:** Video of DIC timelapse images of isolated *Mnemiopsis* cells *in vitro*. An amoebocyte-like granular cell (arrowhead) and highly motile stellate cell (arrow) are visible.

**Supp. Video 2**: Video of DIC timelapse images of an isolated *Mnemiopsis* stellate cell *in vitro*. The cell initially has two large processes, then absorbs them and subsequently produces several additional processes. The cell then begins to crawl out of view.

**Supp. Video 3:** Merged brightfield and fluorescent video of a live *Mnemiopsis* motile, stellate cell that is phagocytosing fluorescent *E. coli* (green). *Mnemiopsis* cells were labeled with Lysotracker Red. Scale bar is 10 µm.

**Supp. Video 4**: Merged brightfield and fluorescent video of a live *Mnemiopsis* motile, stellate cell undergoing ETosis after *in vitro* exposure to pHrodo-*E. coli*. The cell moves into view, retracts its processes, spins, and exudes its nuclear material. DNA (Hoechst, blue), pHrodo-*E. coli* (red)

**Supp. Video 5:** 3D reconstruction of confocal stack of *Mnemiopsis* extracellular DNA traps. DNA (Hoechst, white) is extracellular, in filamentous "nets"; lysosomal marker (Lysotracker, green) denotes cellular debris; bacteria are ensnared in the "nets" (pHrodo-*E. coli*, red)

**Supp. Video 6:** Merged brightfield and fluorescent video of a live *Mnemiopsis* motile, stellate cell that is phagocytosing fluorescent *E. coli* (red). Scale bar is 10 µm.
